# Counterintuitive effect of antiviral therapy on influenza A-SARS-CoV-2 coinfection due to viral interference

**DOI:** 10.1101/2023.02.07.527372

**Authors:** Nagarjuna R. Cheemarla, Valia T. Mihaylova, Timothy A. Watkins, Ellen F. Foxman

## Abstract

The resurgence of influenza and continued circulation of SARS-CoV-2 raise the question of how these viruses interact in a co-exposed host. Here we studied virus-virus and host-virus interactions during influenza A virus (IAV) -SARS-CoV-2 coinfection using differentiated cultures of the human airway epithelium. Coexposure to IAV enhanced the tissue antiviral response during SARS-CoV-2 infection and suppressed SARS-CoV-2 replication. Oseltamivir, an antiviral targeting influenza, reduced IAV replication during coinfection but also reduced the antiviral response and paradoxically restored SARS-CoV-2 replication. These results highlight the importance of diagnosing coinfections and compel further study of how coinfections impact the outcome of antiviral therapy.

## Background

The resurgence of influenza in 2022-23, continued circulation of SARS-CoV-2 variants, and anticipated winter seasonality of both viruses in the future have focused attention on understanding how these viruses interact in a co-exposed host. While intuitively coinfections might be expected to exacerbate disease, increasing evidence suggests that viral interference can also occur, in infection with one virus decreases replication of the other [1-4]. Since antivirals for both SARS-CoV-2 and influenza virus are currently available, in this study we sought to explore the implications of virus-virus interactions for antiviral therapy.

One demonstrated mechanism of interference among respiratory viruses is induction of the interferon response, a broad antiviral defense mechanism that is induced by most respiratory viruses, and also suppresses replication of most viruses[5-8]. Within the airway epithelium, the replicating viral genome is the initiating trigger of the interferon response[9]. Viral RNA is sensed by cytosolic innate immune sensors, leading to expression of interferon stimulated genes (ISGs), which include >300 genes that block viral replication through diverse mechanisms[10]. Viral RNA sensing also leads to secretion of interferons which induce ISG expression in uninfected bystander cells, creating an antiviral state throughout the tissue. Since ISG induction is driven by viral load, there is a possibility that therapeutically blocking replication of the interfering virus could have the paradoxical effect of making the tissue more permissive to infection and increasing replication of the coinfecting virus. Here we used the air-liquid interface (ALI) culture model, a tissue-like in vitro model of the human airway epithelium, to explore virus-virus and virus-host response dynamics and the effect of the influenza inhibitor oseltamivir during IAV-SARS-CoV-2 coinfection.

## Methods

### Primary human bronchial epithelial cells

Low passage primary human bronchial epithelial cells (HBEC) from healthy adult donors were obtained commercially (Lonza, Walkersville, MD, USA) and cultured at air-liquid interface according to the manufacturer’s instructions (Stem Cell Technologies, Vancouver, BC, Canada). Differentiated and mature epithelial cultures displayed beating cilia and mucus production at the time of viral infection. Lonza guarantees that all tissue utilized for human cell products is ethically obtained with donor informed consent in accordance with processes approved by an Institutional Review Board or comparable independent review body.

### Generation of virus stocks

IAV (H1N1 pdm09, strain A/California/07/2009; ATCC VR-1894) was amplified in MDCK cells (MDCK NBL-2, CCL-34, ATCC) and filtered cell lysates were used as virus stocks. IAV stock titers were determined using MDCK cells as described previously (5). SARS-CoV-2 (USA-WA1/2020; BEI Resources) was generously provided by the Wilen laboratory (Yale University, New Haven, CT). To generate virus stocks, SARS-CoV-2 was cultured on Vero E6 cells (CRL-1586; ATCC) and filtered supernatant was used as the virus stock. SARS-CoV-2 titer was determined by plaque assay using Vero E6 cells as described previously [5, 6].

### In vitro infections

Primary human bronchial epithelial cell ALI cultures were inoculated for 1 hr on the apical surface as described previously [5, 6]. For each virus, we used the lowest MOI for which we observed reproducible exponential replication, which was MOI 0.01 for IAV (H1N1pdm09), and MOI 0.1 for SARS-CoV-2 (USA-WA1/2020). For oseltamivir treatment starting at 16 hr post-inoculation, oseltamivir (item no: 15779, Cayman Chemical, Ann Arbor, Michigan, USA) at a final concentration of 1μM was added to the basolateral medium at 16 hr, 40 hr, and 64 hr, and cultures were collected at 72 hr to determine viral load and host response. For treatment starting at 40 hr, oseltamivir was added at 40 and 64 hr only, and cultures were collected at 72hr. All incubations were performed at 35°C to simulate the temperature of the conducting airways.

### RT-qPCR

For RT-qPCR, RNA was isolated from each well of differentiated epithelial cells by incubating each insert with 350 μl lysis buffer at room temperature for 5 min, followed by RNA isolation using the Aurum total RNA isolation mini kit (Bio-Rad, Hercules, CA) and cDNA synthesis using iScript cDNA synthesis kit (Bio-Rad, Hercules, CA). To quantify IAV viral RNA and mRNA levels for ISGs and the housekeeping gene HPRT, RT-qPCR was performed using SYBR green iTaq universal (Bio-Rad) per the manufacturer’s instructions. RT-qPCR for SARS-CoV-2 from cell lysates was performed using the TaqMan assay for the CDC N1 gene primers and probes from Integrated DNA Technologies (catalog no. 10006600; Integrated DNA Technologies) using the Luna Universal Probe One-Step RT-qPCR Kit (New England Biolabs) as previously described (6). Viral RNA per ng total RNA is represented as fold change from the limit of detection (40 cycles of PCR) as 2^40-Ct^. ISG mRNA levels are shown as relative to the level of mRNA for the housekeeping gene HRPT (2^−ΔΔCT^).

For RT-qPCR using SYBR green assay, the following primer sets were used: HPRT (forward: 5′-TGGTCAGGCAGTATAATCCAAAG-3′; reverse: 5′-TTTCAAATCCAACAAAGTCTGGC-3′), MX1 (forward: 5′-AGAGAAGGTGAGAAGCTGATCC-3′; reverse: 5′-TTCTTCCAGCTCCTTCTC TCTG-3′), BST2 (forward: 5′-CACACTGTGATGGCCCTAAT-3′; reverse: 5′-TGTAGTGATCTC TCCCTCAAGC-3′), RSAD2 (forward: TCGCTATCTCCTGTGACAGC; reverse: CACCACCTCCTCAGCTTTTG), ISG 15 (forward: CATCTTTGCCAGTACAGGAGC; reverse: GGGACACCTGGAATTCGTTG), IAV Cal09 (forward: GGGTGGACAGGGATGGTAGA; reverse: TCTGTGTGCTCTTCAGGTCG).

### Statistical analysis

Statistical analyses were performed using GraphPad Prism (version 9.3.1). P value for statistical significance of differences between conditions was determined using the Mann–Whitney test and *p* < 0.05 was considered statistically significant.

## Results

To determine the impact of coinfection on IAV and SARS-CoV-2 replication, we compared single infections to simultaneous and sequential coinfections of IAV (H1N1pdm09) and SARS-CoV-2 (WA/01 strain) at low multiplicity of infection (**Fig 1**). Infection with IAV three days prior to SARS-CoV-2 led to more than 10,000-fold suppression of SARS-CoV-2 replication by 72 hr post-infection (Fig 1B, center) Simultaneous infection with SARS-CoV-2 and IAV also led to a significant decrease in SARS-CoV-2 viral load by day three post infection (**Fig 1B**, right) Conversely, SARS-CoV-2 co-infection under these conditions did not suppress IAV replication (**Fig S1**). Compared to SARS-CoV-2 infection alone, expression of the interferon stimulated genes ISG15 and MX1 were significantly elevated three days post-SARS-CoV-2 infection in sequential and simultaneous coinfections with IAV and in IAV infection alone (**Fig 1C, D**). Taken together, these findings show that IAV infection induces a more robust ISG expression in the human airway epithelium than SARS-CoV-2 in this model, that IAV coinfection enhances the antiviral response during SARS-CoV-2 infection, and that co-infection with IAV restricts SARS-CoV-2 replication.

**Figure 1:**
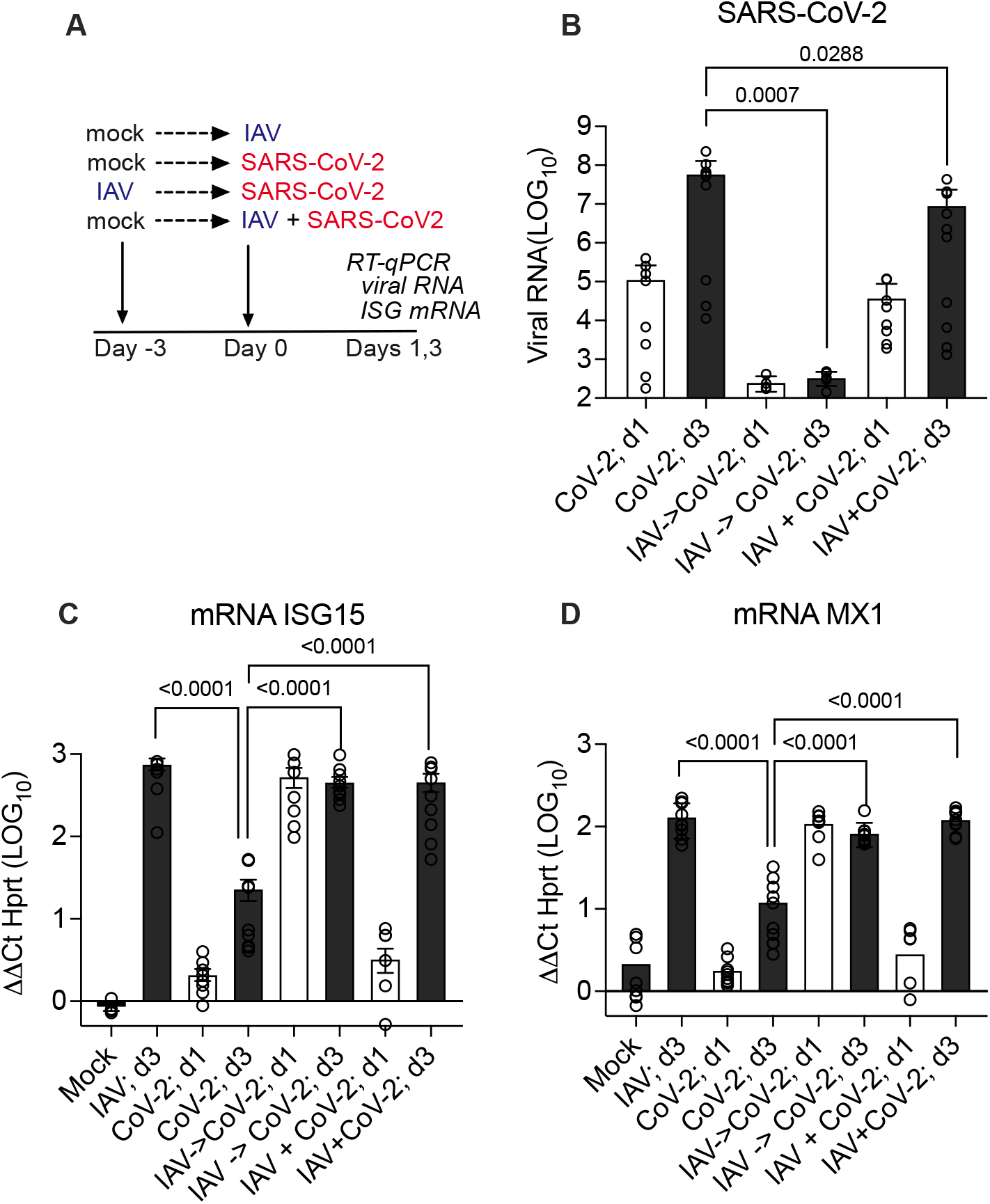
Effect of prior or simultaneous Influenza A virus (IAV) infection on SARS-CoV-2 replication. **(A)** Experimental design of simultaneous or sequential infection in differentiated human airway epithelial cultures. (**B**) SARS-CoV-2 RNA quantification by RT-qPCR on day 1 (24 hr post-CoV-2 infection; white bars) and day 3 (72 hr; grey bars) represented as fold change from detection limit. (**C, D**) MRNA level of interferon stimulated genes ISG15 or MX1 by RT-qPCR on day 1 (24 hr post-CoV-2 infection; white bars) and day 3 (72 hr; grey bars) relative to mRNA level of housekeeping gene HPRT. Graphs show combined resuls of two independent experiments using primary human bronchial epithelial cultures from different healthy adult donors, each with 4-5 replicates per condition. Mean and S.E.M. of 9-10 replicates is shown. Mann-Whitney p-values are shown for conditions that differ significantly from SARS-CoV-2 infection only on day 3.

Since we observed suppression of SARS-CoV-2 replication by IAV but not vice versa, we next asked how treatment with oseltamivir, an FDA-approved influenza virus replication inhibitor, impacts IAV and SARS-CoV-2 replication during co-infection. ALI cultures were infected with IAV, SARS-CoV-2, or both viruses and treated with oseltamivir starting at 15 hours post infection (**Fig 2A**). As expected, oseltamivir treatment during IAV single infection or coinfection significantly reduced IAV viral load by day 3 post infection (**Fig 2B**). While oseltamivir alone had no effect on SARS-CoV-2 replication (**Fig S2**), oseltamivir treatment rescued SARS-CoV-2 replication during simultaneous infection (**Fig 2C**). Further analysis of how the timing of oseltamivir impacts IAV replication and host-virus interactions showed that when administered at 16 hr post inoculation, oseltamivir significantly decreased both IAV replication and host tissue ISG expression and viral load by 72 hr, but when administered at 40 hr post-infection, treatment did not impact viral replication or ISG levels at 72 hr (**Fig S3**) Taken together, these data confirm that IAV co-infection suppresses SARS-CoV-2 replication and indicate that blocking IAV replication with oseltamivir releases SARS-CoV-2 from IAV-mediated interference.

**Figure 2:**
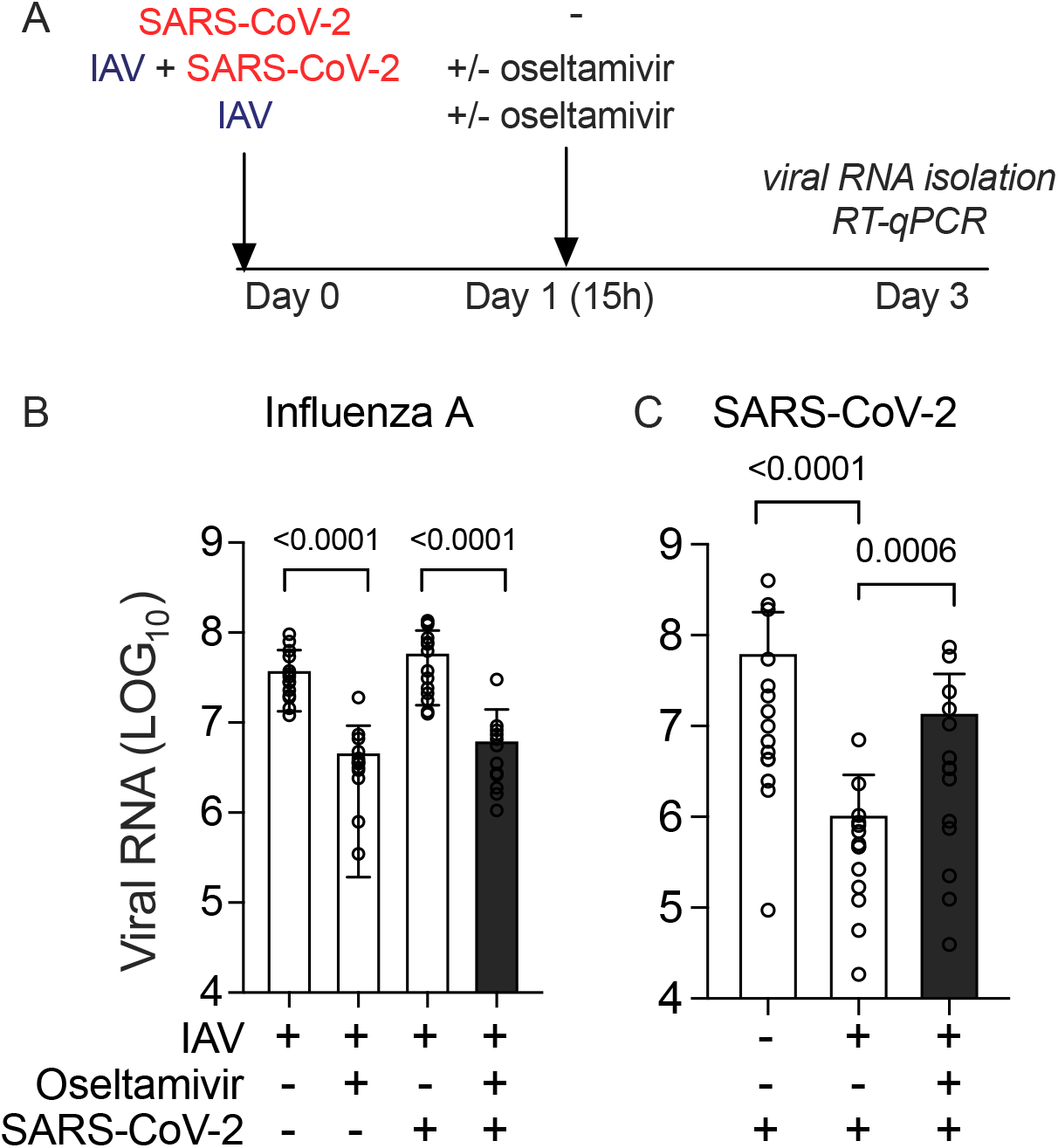
Effect of oseltamivir on influenza A and SARS-CoV-2 replication during simultaneous infection. **(A)** Experimental design. (**B, C**) Quantification of IAV (B) and SARS-CoV-2 viral RNA (C) by RT-qPCR at days 3 post single infection or simultaneous coinfection, with or without addition of oseltamivir starting at 15 hr. Viral RNA is expressed as fold change from detection limit. Bars show mean and S.E.M. of 14 replicates, representing pooled data from three independent experiments, each with 4-6 replicates per condition. Conditions with coinfection and oseltamivir are highlighted with shaded bar. Mann-Whitney p-values are shown.

## Discussion

Circulation of influenza virus reached a historically low level during the first two years of the COVID-19 pandemic, however there has been striking resurgence of influenza during the 2022-23 winter season as pandemic mitigation measures ease, increasing the likelihood of influenza-SARS-CoV-2 coinfections [11]. A better understanding of how these viruses interact with each other and with the host in co-exposed subjects is crucial to inform clinical practice, both now and in the future, since both viruses are likely to continue to co-circulate with winter seasonality typical of influenza viruses and coronaviruses.

The range of clinical outcomes following influenza and SARS-CoV-2 co-exposure remains unclear. A large study of viral coinfections in hospitalized patients found increased clinical severity metrics in subjects with influenza-SARS-CoV-2 coinfection, but this study had potential observation biases in that only hospitalized patients were observed, and that subjects with more severe disease were more likely to undergo multiplex virology testing[12].

Experimental models show varied outcomes. Coinfection in a ferret model and in the highly SARS-CoV-2 susceptible K18-hACE2 transgenic mouse showed increased disease severity during coinfection compared to single infections [13, 14]. In contrast, viral interference was seen in the Syrian golden hamster model, with one study showing IAV suppressing SARS-CoV-2 replication, and other showing SARS-CoV-2 suppression of IAV [1, 3]. Studies using differentiated human airway epithelial cultures, like this study, have reported interference of influenza with SARS-CoV-2, with the degree of interference varying with virus strains and timing of infections [2, 4]. For studies showing interference, the host interferon response appears to be an important mechanism, since the kinetics and magnitude of ISG expression predict interference, and blocking host cell signaling pathways required for this response rescues replication of the suppressed virus [1-4]. Interferon response-dependent interference was also described in previous work from our group and others showing suppression of both influenza A virus and SARS-CoV-2 by prior rhinovirus infection [5-7, 15]. Together these results suggest that many factors influence whether IAV and SARS-CoV-2 will show interference or potentiation during coinfection, and that host susceptibility, as well as the relative timing, infectious dose, and virus strains involved, likely all play important roles in determining the outcome.

The airway epithelial cultures used in this study model the earliest stages of infection in a target tissue with robust innate antiviral immune responses: the human airway epithelium. In the model used here, IAV causes a mild infection with minimal tissue damage that is well-controlled by the innate immune response[5]. This model probably best represents a mild upper respiratory tract infection, and we speculate that mild, well-controlled infections would be the most likely to result in interference, rather than potentiation, of a subsequent or concurrent viral exposure. A limitation of this model is the lack of other cell types present in the respiratory mucosa in human subjects, including resident and recruited leukocytes, which can also contribute to the innate (and adaptive) immune response. In a prior study, we examined the nasal interferon response in subjects with mild-to-moderate SARS-CoV-2 infection and we observed a robust and long-lasting nasal interferon response in SARS-Cov-2 infected subjects, with nasal ISG elevation lasting up to four weeks post-infection [6]. These data suggest that the ALI culture model may underestimate the potential for SARS-CoV-2 to interfere with subsequent viral infections in human subjects.

While the extent to which viral interference operates during coinfections in human subjects is still unknown, if this does occur, this study indicates important implications for therapy. Rapid point-of-care tests and antiviral drugs are available for both influenza A and SARS-CoV-2. While multiplex virology testing has gained traction over the past several years, it is still common practice for patients to undergo testing for one virus at a time and receive therapy if the test is positive. However, the results presented here indicate that in the case of an undiagnosed coinfection, treating only one virus could have the unanticipated consequence of increasing replication of the other. A clinical implication is that even when a point of care test for one of these viruses is positive, testing for coinfection may be advisable prior to treatment.

Further studies in both animal models and human subjects are needed to fully understand the range of IAV-SARS-CoV-2 coinfection outcomes. However, this study contributes to growing evidence of the potential for interference between these viruses, illustrates that antiviral therapy during coinfection can have the unexpected outcome of promoting infection with the un-targeted virus due to viral interference. As seasonal respiratory viruses increase in prevalence to pre-pandemic levels, these results highlight the importance of diagnosing coinfections and compel further study of how coinfections impact the outcome of antiviral therapy.

## Acknowledgements and Funding

## Acknowledgements

We thank the Wilen laboratory for providing the SARS-CoV-2 isolate and VeroE6 cells.

## Funding Sources

Funding was provided by Fast Grants for COVID-19 from Emergent Ventures at the Mercatus Center of George Mason University (E.F. Foxman); the Yale Department of Laboratory Medicine and COVID-19 Dean’s Fund (E.F. Foxman); the Gruber Foundation Fellowship (T.A. Watkins); and the National Institutes of Health (grant T32AI007019 to T.A. Watkins).

## Declaration of interests

All authors declare no conflicts of interest.

## Author contributions

Conceptualization: NRC and EFF; Investigation: NRC, VTM, and TAW; Writing: Original Draft preparation: NRC and EFF, Reviewing and Editing: all authors; Data visualization: NRC and EFF; Supervision: EFF, Acquisition of funding: EFF

## Notes

### Competing Interest Statement

The authors have declared no competing interest.

## REFERENCES CITED

1. Halfmann PJ, Nakajima N, Sato Y, et al. SARS-CoV-2 Interference of Influenza Virus Replication in Syrian Hamsters. The Journal of infectious diseases 2022; 225:282–6.

2. Essaidi-Laziosi MP, Alvarez CM, Puhach OP, et al. Sequential infections with rhinovirus and influenza modulate the replicative capacity of SARS-CoV-2 in the upper respiratory tract. Emerg Microbes Infect 2021:1–26.

3. Oishi K, Horiuchi S, Minkoff JM, tenOever BR. The Host Response to Influenza A Virus Interferes with SARS-CoV-2 Replication during Coinfection. J Virol 2022; 96:e0076522.

4. Dee K, Schultz V, Haney J, Bissett LA, Magill C, Murcia PR. Influenza A and respiratory syncytial virus trigger a cellular response that blocks severe acute respiratory syndrome virus 2 infection in the respiratory tract. The Journal of infectious diseases 2022.

5. Wu A, Mihaylova VT, Landry ML, Foxman EF. Interference between rhinovirus and influenza A virus: a clinical data analysis and experimental infection study. Lancet Microbe 2020; 1:e254–e62.

6. Cheemarla NR, Watkins TA, Mihaylova VT, et al. Dynamic innate immune response determines susceptibility to SARS-CoV-2 infection and early replication kinetics. J Exp Med 2021; 218.

7. Gonzalez AJ, Ijezie EC, Balemba OB, Miura TA. Attenuation of Influenza A Virus Disease Severity by Viral Coinfection in a Mouse Model. J Virol 2018; 92.

8. Cox G, Gonzalez AJ, Ijezie EC, et al. Priming With Rhinovirus Protects Mice Against a Lethal Pulmonary Coronavirus Infection. Front Immunol 2022; 13:886611.

9. Park A, Iwasaki A. Type I and Type III Interferons - Induction, Signaling, Evasion, and Application to Combat COVID-19. Cell Host Microbe 2020; 27:870–8.

10. Schneider WM, Chevillotte MD, Rice CM. Interferon-stimulated genes: a complex web of host defenses. Annual review of immunology 2014; 32:513–45.

11. World Health Organization: FluNet. Available at: https://www.who.int/tools/flunet.

12. Swets MC, Russell CD, Harrison EM, et al. SARS-CoV-2 co-infection with influenza viruses, respiratory syncytial virus, or adenoviruses. Lancet 2022; 399:1463–4.

13. Kim EH, Nguyen TQ, Casel MAB, et al. Coinfection with SARS-CoV-2 and Influenza A Virus Increases Disease Severity and Impairs Neutralizing Antibody and CD4(+) T Cell Responses. J Virol 2022; 96:e0187321.

14. Huang Y, Skarlupka AL, Jang H, et al. SARS-CoV-2 and Influenza A Virus Coinfections in Ferrets. J Virol 2022; 96:e0179121.

15. Dee K, Goldfarb DM, Haney J, et al. Human Rhinovirus Infection Blocks Severe Acute Respiratory Syndrome Coronavirus 2 Replication Within the Respiratory Epithelium: Implications for COVID-19 Epidemiology. The Journal of infectious diseases 2021; 224:31–8.

