## Supplemental Figures for "Counterintuitive effect of antiviral therapy on influenza A-SARS-CoV-2 coinfection due to viral interference"

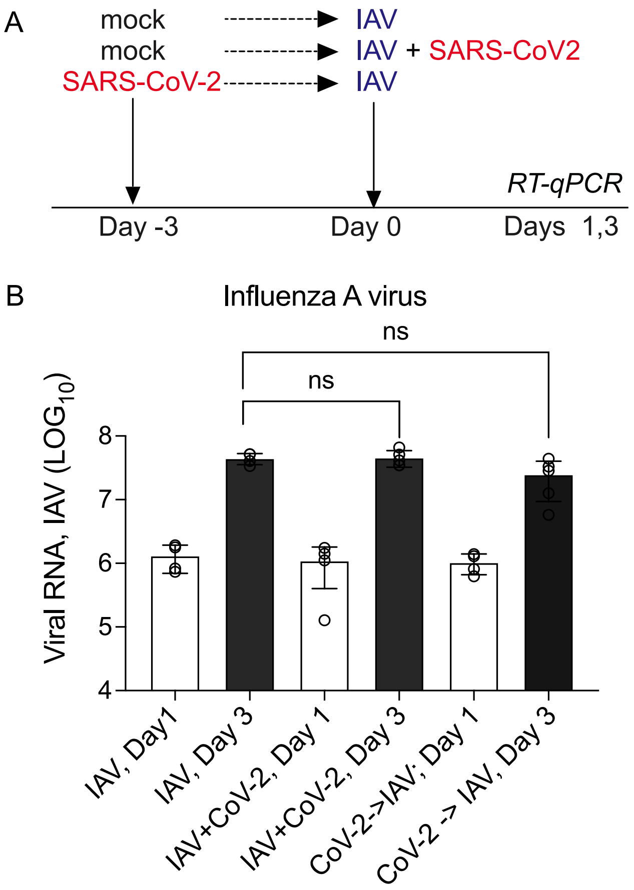


**Fig S1 (related to Fig 1). Effect of SARS-CoV-2 on influenza A virus replication.** (A) Experimental design of simultaneous or sequential infection in differentiated human airway epithelial cultures. (B) IAV (H1N1pdm09) RNA quantification by RT-qPCR on days 1 and 3 represented as fold change from detection limit. Mean and S.E.M. of 5 replicates per condition is shown.

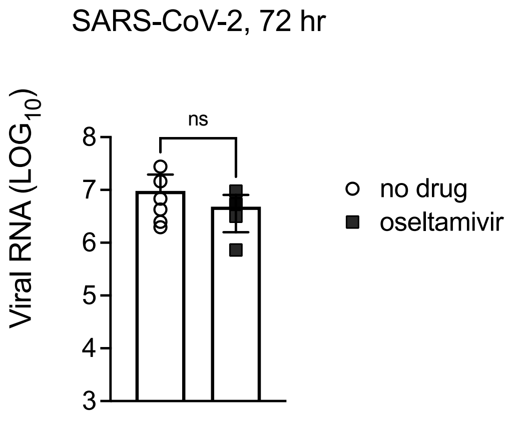


**Fig S2 (related to Fig 2). Effect of Oseltamivir on SARS-CoV-2 replication.** SARS-CoV-2 infected cutlures were treated with oseltamivir starting at 15 hr as in Fig 2 and collected at 72 hr for viral RNA isolation and RT-qPCR for the SARS-CoV-2 N1 gene. Mean and S.D. of 6 replicates per condition are shown.

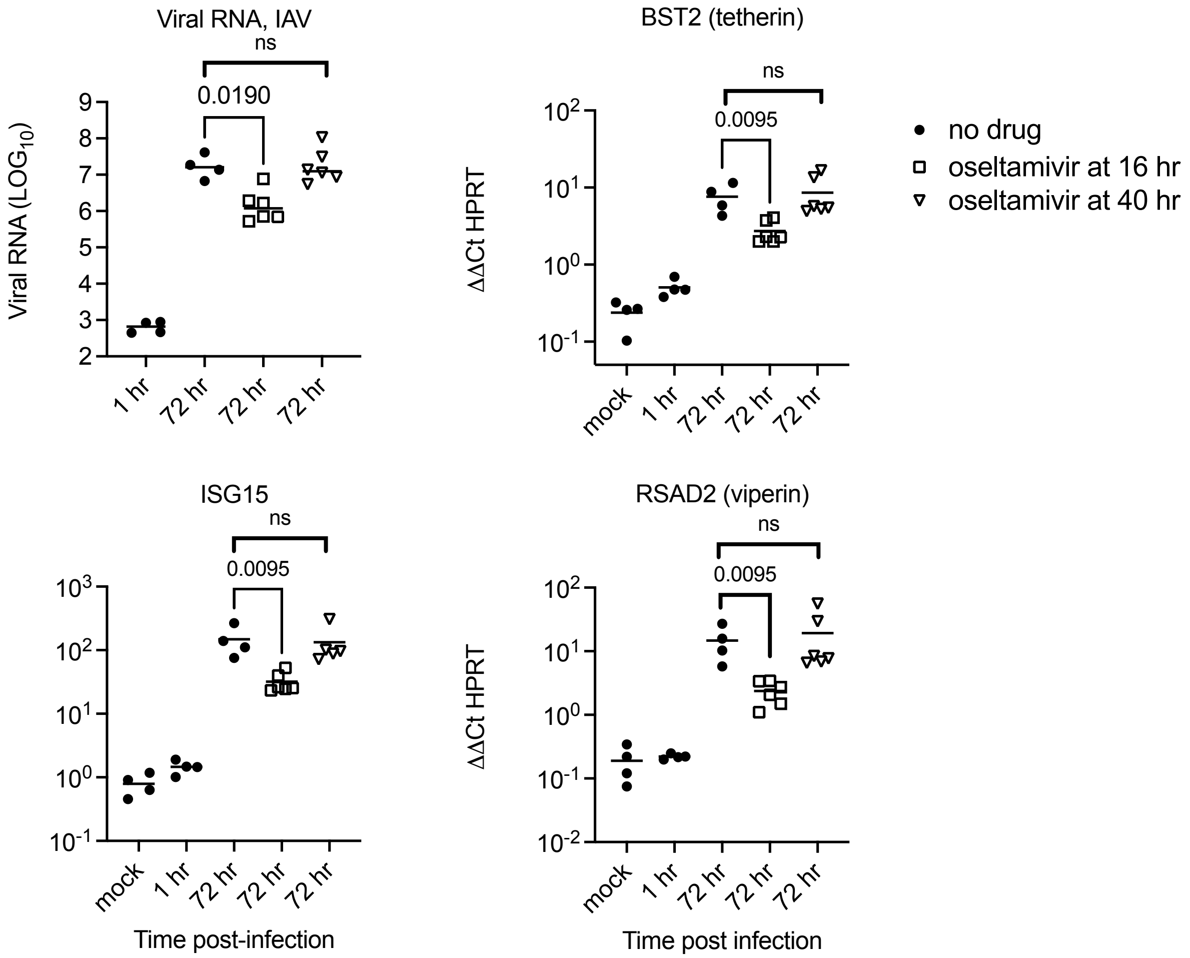


**Fig S3 (related to Figure 2). Effect of timing of Oseltamivir on influenza A replication and ISG induction.** ALI cultures were infected with IAV H1N1pdm09 and oseltamivir was added to the basolateral media starting 16 hr or 40 hr post-inoculation. RNA isolation and RT-qPCR for ISGs were performed at 72 hr post-infection. (A)IAV RNA level at 72 hr post infection in RNA isolated from ALI culture tissue, plotted as fold change from limit of detection. (B-D) ISG mRNA levels shown relative to mRNA for HPRT. Mean and S.D. of 4-5 replicated per conidition is shown. Mann-Whitney p-values comparing 72 hr timepoint with and without oseltamivir are indicated.
